# Myxobacteria isolated from RAS: Ecology and significance as off-flavor producers

**DOI:** 10.1101/2025.04.10.648225

**Authors:** Julia Södergren, Pedro Martínez Noguera, Mikael Agerlin Pedersen, Niels O. G. Jørgensen, Raju Podduturi, Mette H. Nicolaisen

## Abstract

Despite advances in the operation of recirculating aquaculture systems (RAS), accumulation of earthy-muddy off-flavors in the fish remains a potential risk. Myxobacteria (Myxococcota) are reported to be among the most abundant geosmin synthase-harboring groups in RAS, but previous isolation attempts have been unsuccessful, limiting the knowledge of their role in off-flavor production. For the first time, we successfully isolated two geosmin-producing myxobacteria from RAS: *Myxococcus virescens* AT3 and *Corallococcus exiguus* AT4. Cell-specific geosmin production varied with the nutrient content in different media but was highest in a low-nutrient medium and when cultivated in water from RAS. Cultivation in RAS water also stimulated the production of other volatile organic compounds (VOCs). Newly identified potential off-flavor compounds included 4-methyl-2-heptanone (“forest” odor), 3-methyl-1-butanol (“medicinal” and “chemical”), and a presumptive sesquiterpenoid described as “musty,” “earthy,” and “flowery.” The previously known off-flavor compound dimethyl sulfide was also detected. Myxobacteria have previously been proposed as keystone bacteria in the environment due to their predatory lifestyle. In predation assays using isolated bacteria from RAS, *M. virescens* AT3 and *C. exiguus* AT4 could successfully feed on 15 of 16 tested strains, suggesting a large influence on the biology of RAS microbiomes. The combination of predatory behavior and potent production of geosmin and other VOCs underscores the ecological and sensory impact of these bacteria in RAS. Understanding their behavior and metabolic outputs is critical to developing strategies for mitigating off-flavors in RAS.

**IMPORTANCE:** Issues with off-flavored fish in recirculating aquaculture systems (RAS) due to the presence of the earthy-musty smelling compounds geosmin and 2-MIB is considered one of the industry’s most economically significant challenges. Knowledge of conditions that affect off-flavor production is essential information in the development of viable solutions for its mitigation. Little is known about the function of these microbially produced compounds or the conditions that trigger their production, especially in the underexplored myxobacteria. Investigation of natural isolates is crucial to determine the function of the genes involved and their differential expression in response to environmental cues. While myxobacteria in RAS have been previously shown to harbor the geosmin synthase gene through molecular studies, the present study is the first attempt to isolate these bacteria from RAS and quantify their geosmin production under different nutrient conditions. Through cultivation-based methods, we demonstrate their production of both known and novel compounds with earthy attributes.

## INTRODUCTION

The industry of land-based fish farming using recirculating aquaculture systems (RAS) has since its beginning experienced problems with off-flavored fish, mainly attributed to the presence of earthy-musty smelling compounds such as geosmin and 2-methylisoborneol (2-MIB) in the fish flesh (Abd El-Hack et al., 2022). The off-flavors make up a costly problem, which is currently only resolved by the fish undergoing a depuration process at the end of the growth cycle, entailing increased water usage and additional manual labor which reduces the farmers’ profit. Moreover, the animal welfare is challenged due to the lack of feeding during the depuration (Tucker, 2000).

In an attempt to better understand the processes controlling the off-flavor issues, increased biological knowledge about the production of geosmin and 2-MIB have been sought. These two compounds have historically been known to be produced by Cyanobacteria, Actinobacteria (mainly the genus *Streptomyces*), and myxobacteria (Myxococcota). While only Cyanobacteria and *Streptomyces* have previously been isolated, DNA sequencing shows that all three groups occur in RAS, with Cyanobacteria being most important in outdoor systems due to the need of light for photosynthesis (Lukassen et al., 2017). While *Streptomyces* have been assumed main producer of geosmin in RAS (Auffret et al., 2011; Schrader, 2013; Schrader & Summerfelt, 2010), a recent study points to myxobacteria being the most abundant of the three groups in 26 European RAS, at least in terms of copies of the geosmin synthase gene *geoA* (Lukassen et al., 2022). Myxobacteria have mainly been studied for their particular, multicellular lifestyle, and for being prolific producers of a plethora of bioactive compounds and have received less focus as geosmin producers than *Streptomyces.* Isolation of myxobacteria is complicated due to their notoriously slow growth, often in the shape of thin, transparent swarms, where contamination of faster growing species is common (Shimkets et al., 2006). While it has been known for over 40 years that myxobacteria indeed produce geosmin (Trowitzsch et al., 1981), failed attempts to isolate them from RAS is likely the reason for the misconception that *Streptomyces* would be the main geosmin producer in indoor RAS facilities. Interestingly, while *Streptomyces* has been frequently isolated from RAS, presence of copies of the *geoA* gene of *Streptomyces* origin was below limit of the detection in gene libraries of five RAS, located in Denmark and Scotland, with geosmin levels of 100-650 ng L^-1^ (Lukassen et al., 2017).

Myxobacteria is the conventional name for the Gram-negative bacteria belonging to the phylum Myxococcota. The name “myxo” derives from the Greek word “muxa” which means slime, due to the bacteria growing in a gliding fashion on surfaces in slime sheets. Myxobacteria are highly social bacteria, which both move and feed cooperatively in groups through predation on other bacteria. This is a unique behavior to myxobacteria. The predation occurs when individual cells together have produced a sufficient amount of bioactive compounds that can lyse the prey cells (Contreras-Moreno et al., 2024). The bioactive compounds include hydrolases, antibiotics and additional secondary metabolites. Upon lysis, the myxobacteria consume the hydrolyzed cell material. Another unique phenotypic character in myxobacteria is the formation of so-called fruiting bodies, which are three-dimensional mounds of aggregated dead and living myxobacteria that are formed when nutrients are scarce. The fruiting bodies serve as a survival tactic, protecting the bacteria and myxospores inside the fruiting body from desiccation, high temperatures and other environmental threats until living conditions are once again better, allowing the spores to germinate (Muñoz-Dorado et al., 2016).

Due to their predatory and saprophytic lifestyle, myxobacteria have previously been acknowledged as important contributors to nutrient cycling and shapers of the microbiome of their habitat. Myxobacteria have been shown to represent around 2% of the total bacterial operational taxonomic units (OTUs) in the Earth Microbiome Project, and they are distributed around the globe in saline and non-saline soils, sediments and water, making this group one of the most widespread and diverse phyla on our planet (J. Wang et al., 2021). Myxobacteria are reported to make up 60% of bacterivores identified in 11 European organic and mineral soils from different climatic zones, making them dominant in numbers compared to the traditionally considered eukaryotic micropredators (Petters et al., 2021). The high abundance of myxobacterial bacterivores is speculated to characterize them as keystone taxa that exert a disproportionate influence on the structure of the microbial community they inhabit (Li et al., 2024; Petters et al., 2021)

Whether the ability of myxobacteria to predate other bacteria and produce geosmin is linked, has recently been documented. Two different studies of the transcriptomic changes when *Myxococcus xanthus* encounters prey show an upregulation of the genes involved in geosmin production (Pérez et al., 2022; C. Wang et al., 2023), suggesting that this volatile compound provides some advantage to predation. In a study of the influence of geosmin on the behavior of the nematode *Caenorhabditis elegans*, it was found that geosmin repels the bacterivorous nematodes (Zaroubi et al., 2022), supporting that geosmin could bring some benefits when competing for prey.

Historically, geosmin and 2-MIB have been determined as the main culprits in earthy-tasting fish, but at least 10 other compounds with “earthy” sensory attributes have also been identified in RAS-reared fish (Noguera et al., 2024). To help the RAS industry solve the potential problem with off-flavored fish, these other potential contributing compounds should not be overlooked when analyzing off-flavor issues. Since myxobacteria are known to produce a range of terpenoids and pyrazines (Schulz & Dickschat, 2007; Yamada et al., 2015), their presence in RAS facilities could likely contribute to the generation of other off-flavors, beyond geosmin.

Here, we present for the first time geosmin-producing myxobacteria isolated from RAS, comparing the geosmin production of the two isolates and examining how the production depends on nutritional factors, as well as an identification and description of other potential off-flavors that they produce. Furthermore, the potential ecological impact of myxobacteria on other bacteria in the system is investigated in a predation assay on selected RAS bacteria.

## MATERIAL AND METHODS

### Isolation procedure of myxobacteria and general RAS bacteria

Water, biofilm and biomedia samples were collected in screwcap bottles from two outdoor RAS systems for production of rainbow trout in Denmark. The bottles were rigorously shaken to release solid material from biomedia and biofilms. The isolation of myxobacteria from these samples was performed by two baiting techniques. Petri dishes were prepared with RAS water agar (WA), made with autoclaved water collected from the RAS tanks and 1.5% (w/v) agar, supplemented with 30 mg/ml of cycloheximide to suppress fungal growth. To bait myxobacteria, either a cross streak of *E. coli* was placed on top of the agar surface, or sterile rabbit dung pellets were partly submerged into the agar before it had fully solidified (L. P. Zhang et al., 2003). Solid material from the sample bottles was inoculated at the edge of the cross streak, and adjacent to the dung pellets. The plates were incubated at 30°C and examined regularly for the appearance of fruiting bodies. When fruiting bodies appeared, the culture was transferred by dabbing the top of the fruiting body with the tip of a sterile needle before inoculating onto VY/2 agar (Bakers’ yeast 0.5% (w/v), CaCl_2_ · 2H_2_O 0.1% (w/v), Cyanocobalamin 0.5 mg/µl, agar 1.5% (w/v)). If the growth of the transferred culture was contaminated with other organisms, repeated transfers from the fruiting body or the swarm edge of the colony were made until the culture was pure.

To determine whether the isolated myxobacteria could prey on other bacteria in the RAS system, bacteria in RAS water were isolated on Petri dishes containing tryptic soy agar (TSA) (Merck KGaA, Darmstadt, Germany), Luria Bertani agar (LBA) (10 g/L tryptone, 10 g/L NaCl, 5 g/L yeast extract, 15 g/L agar), VY/2 agar and Reasoner’s 2A agar (R2A) (Alpha Biosciences, Baltimore, USA) in full and 1/10 concentration, as well as WA prepared as above. Cycloheximide at a final concentration of 30 mg/ml was supplemented to the Petri dishes containing WA, VY/2 and R2A in 1/10 concentration. Individual colonies occurring on the Petri dishes were transferred to new identical medium to test of purity. Finally, the cultivated bacteria were stored in 15% glycerol at -70°C until further analysis. The collection of the bacteria was expected to represent a “biobank of general bacteria” from the RAS facilities.

### Preparation of gDNA, sequencing and phylogenetic analyses

DNA for genome sequencing of the myxobacterial isolates was obtained using phenol-chloroform extraction according to a basic protocol (Wilson, 2001). The whole genome sequencing was performed by Plasmidsaurus using Oxford Nanopore Technology with custom analysis and annotation. The obtained genomes were used for orthologous Average Nucleotide Identity (ANI) analysis with OAT v. 0.93.1 (Lee et al., 2016). Reference genomes for other *Myxococcus* and *Corallococcus* were obtained from NCBI Genbank with accession numbers: PRJNA331492 (*M. fulvus* DSM 16525), PRJNA606434 (*M. vastator* AM301), PRJEB15777 (*M. virescens* DSM 2260), PRJNA1421 (*M. xanthus* DK 1622), PRJNA82779 (*C. coralloides* DSM 2259), PRJNA702492 (*C. exiguus* NCCRE002), PRJNA490141 (*C. interemptor* AB047A), and PRJNA490141 (*C. terminator* CA054A).

DNA was extracted from the biobank of general bacteria using a boiling extraction method. The strains were cultivated in suitable liquid media. One ml of each actively growing strain was transferred to an Eppendorf tube which was boiled for 10 minutes with shaking at 300 rpm on a heat block. The extracted DNA was subsequently used as template in a PCR reaction targeting the 16S rRNA gene with universal primers 27F and 1492R (Lane, 1991). The PCR products were sequenced by Sanger sequencing performed by Eurofins Genomics Europe. A phylogenetic tree was constructed using the obtained 16S rRNA sequences from the RAS biobank rooted on the 16S rRNA sequences of the two myxobacterial isolates. The tree was constructed by first aligning the sequences with ClustalX 2.1, followed by the construction of a neighbor-joining tree with bootstrap values based on 1000 resamplings using MEGA11.

### Predation assay of RAS bacteria

The predation assay was based on a previously described procedure by Berleman and Kirby (2007), with minor modifications. Cells of the myxobacteria and potential prey bacteria from the RAS biobank were harvested mid-log phase and washed twice with TPM buffer (10 mM Tris-HCl (pH 7.6), 1 mM KH_2_PO_4_, 8 mM MgSO_4_). The bacterial pellets were resuspended to an OD_600_ of ∼1, and 2 µl each of myxobacteria and prey were together spotted 0.5 cm apart on individual CFL agar plates (10 mM MOPS (pH 7.6), 1 mM KH_2_PO_4_, 8 mM MgSO_4_, 0.02% (NH_4_)_2_SO_4_, 0.02% citrate, 0.02% pyruvate, 0.1 g/L Casitone, 15 g/L agar). The plate cultures were incubated at 30°C and monitored daily with a stereo dissection microscope for four days, and once more after seven days of incubation. Predation was assessed by clearing of the prey colony. Inhibition of the predator was assessed by a difference in radial expansion of the predator swarm.

### Cultivation conditions for volatile production of isolated myxobacteria

To evaluate the dependence of geosmin production on different nutrients, the isolated myxobacteria were cultivated in four liquid media supporting myxobacterial growth (Table 1) (Shimkets et al., 2006). Geosmin production was also evaluated by cultivation in autoclaved RAS rearing water (RW) from a trout farm. Each of the isolated bacteria was inoculated into 50 ml Falcon tubes containing 20 ml of the different media in replicates of five and incubated obliquely on a rotary shaker (150 rpm) at 30°C for 48 hours.

**Table 1.** Composition of different media used for investigation of volatile production.

| *Medium* |  |  |
| --- | --- | --- |
| VY/2 (complex medium) | Bakers' yeast (commercial yeast cake) | 0.5% (w/v) |
|  | CaCl<sub>2</sub> 2H<sub>2</sub>O | 0.1% (w/v) |
|  | Cyanocobalamin | 0.5 mg/μl |
| CTT (complex medium) | Bacto Casitone | 1% (w/v) |
|  | Tris-HCl | 10 mM |
|  | KH<sub>2</sub>PO<sub>4</sub>-KHPO<sub>4</sub> | 1 mM |
|  | MgSO<sub>4</sub> | 8 mM |
| A1 (defined minimal medium) | Tris-HCl | 1 mM |
|  | KH<sub>2</sub>PO<sub>4</sub>-KHPO<sub>4</sub> | 1 mM |
|  | MgSO<sub>4</sub> | 8 mM |
|  | (NH<sub>4</sub>)<sub>2</sub>SO<sub>4</sub> | 0.5 mg/ml |
|  | L-Asp | 100 μg/ml |
|  | L-Ile | 100 μg/ml |
|  | L-Phe | 100 μg/ml |
|  | L-Leu | 50 μg/ml |
|  | L-Met | 10 μg/ml |
|  | Sodium pyruvate | 0.5% |
|  | Potassium Aspartate | 0.5% |
|  | FeCl<sub>3</sub> | 10 μM |
|  | CaCl<sub>2</sub> | 10 μM |
|  | Cyanocobalamin | 1 μg/ml |
| TPM (starvation medium) | Tris-HCl | 10 mM |
|  | KH<sub>2</sub>PO<sub>4</sub> | 1 mM |
|  | MgSO<sub>4</sub> | 8 mM |

### VOC extraction with Stir-Bar Sorptive Extraction (SBSE)

After two days of incubation, liquid cultures from the VY/2, CTT, A1, TPM and RW media were centrifuged at 4200 x g for 10 min. The supernatant from each of the cultures was used for the extraction of bacterial volatiles. For GC-MS analysis, geosmin and other VOCs in the supernatants were extracted in quintuplicate using a method adapted from Podduturi et al. (2020). A control without inoculum for each medium was also extracted and analyzed by GC-MS. In brief, for the extraction of VOCs and geosmin, a commercial stir bar (Twister®; Gerstel GmbH & Co. KG, Mülheim an der Ruhr, Germany) coated with polydimethylsiloxane (PDMS; length 2 cm and thickness of 1 mm) was added to 20 mL of the liquid samples in a 50-mL glass vial and covered with a plastic lid. The extraction was carried out at room temperature by stirring the twister at 1000 rpm for 120 min. After extraction, the stir bar was washed with cold distilled water, dried with a non-linting tissue, and transferred to thermal desorption tubes.

### Cell quantification through qPCR

The myxobacterial pellet remaining after cultivation for extraction of VOCs was extracted with Genomics Mini AX Bacteria kit (A&A Biotechnology, Gdansk, Poland). The extracted DNA was used to quantify the number of cells in the liquid cultures by qPCR targeting the 16S rRNA gene, using the primers 907F (5’-AAA-CTC-AAA-GGA-ATT-GAC-GG-3’) and 1492R (5’-TAC-GGT-TAC-CTT-GTT-ACG-ACT-T-3’). In brief, 10 µl of Brilliant III Probe Master Mix with ROX (Agilent Technologies), 0.8 µl of each primer, 1 µl of BSA (20 mg/ml), 5.4 µl sterile MilliQ water and 2 µl of template DNA was used in each reaction. The qPCR reaction was performed with an initial denaturation at 95°C for three minutes, followed by 40 cycles of 95°C for 20 seconds, annealing at 58°C for 30 seconds, and a final extension at 95°C for one minute. When quantifying cell numbers from the 16S copies, the number of 16S rRNA gene copies in the respective myxobacterial genomes was taken into account.

### VOC identification through GC-MS analysis

VOCs were desorbed from the twisters through a two-step procedure using an automatic thermal desorption unit (TurboMatrix 350, Perkin Elmer, Shelton, USA). First, primary desorption was carried out by heating the twister to 240°C for 15 min with a carrier gas flow of 50 ml H_2_·min^-1^. Second, the volatiles desorbed were trapped in a Tenax TA trap held at 1°C and then rapidly heated to 280°C for 4 min to complete the secondary desorption. This results in a quick transfer of volatiles from the thermal desorption unit to a gas chromatograph-mass spectrometer (8890, GC-system, coupled with a 5977B MSD from Agilent Technologies, Palo Alto, California) through a temperature-controlled transfer line held at 225°C. A ZB-WAX capillary column (30 m x 0.25 mm x 0.5 µm) was used for the separation of the transferred volatiles using H_2_ as carrier gas at an initial flow rate of 1.4 ml·min^-1^. The GC oven program was set as follows: isothermal the first 10 min at 35°C, then raised to 240°C at the rate of 8°C/min followed by a holding period of 10 min. Mass spectra of the separated volatile compounds were generated after standard electron ionization conditions (70 eV) and detected through QMS, where the mass acquisition settings were set in a hybrid mode combining SCAN and SIM modes. The SCAN mode scanned *m/z* values ranging from 15 to 300, while the selected ion monitoring mode (SIM) specifically targeted *m/z* 95 and 107 during the first 25 min (for 2-methylisoborneol) and *m/z* 112 (for geosmin) for enhanced sensitivity. SIM chromatograms were processed using the MSD ChemStation software (v.E.02.00, Agilent Technologies). To analyze the GC-MS data in an untargeted way (to obtain volatile profiles), peak areas and mass spectra were extracted from the SCAN chromatograms using the PARAFAC2 algorithm-based software PARADISe (Quintanilla-Casas et al., 2023).

For geosmin quantification in the liquid cultures, external calibration curves were prepared in duplicate from a GC-grade mixture solution (1:1 v/v) of geosmin and 2-MIB in a dilution series of 10, 50, 100, 250, 500, 1,000, 5,000, 10,000 and 25,000 ng·L^-1^ in deionized water.

For the untargeted GC-MS analyses, control samples were used to identify volatiles associated with the media. In this way microbial VOCs could be selected in the remaining samples. Peak areas were later again normalized to the number of counted cells to decouple volatile production from bacterial growth (VOC production/bacterial cell).

### Screening of odor-active compounds by GC-O analysis

VOCs were extracted from one of the strains grown in the complex CTT medium by SBSE and desorbed using the method previously described. For GC-olfactometry (GC-O), a gas chromatograph-mass spectrometer (7890B, GC-system with a Low Thermal Mass DB-WAX module, coupled with a 5977B MSD from Agilent Technologies, Palo Alto, California) and an olfactory detection port (ODP2, Gerstel GmbH & Co., Germany) were used. Flow between the MSD and ODP was split at the splitter in a ratio of 1.7:5.8. A DB-WAX capillary column (30 m x 0.25 mm x 0.5 µm) was used for the separation of the transferred volatiles using H_2_ as carrier gas at an initial flow rate of 1.677 ml·min^-1^. The GC oven program was set as previously described. MS detection was obtained by electron ionization mode at 70 eV with a scanning range of 15–300 m/z. During the analysis, the ODP was constantly supplied with humidified air.

A panel of 5 judges (3 males/2 females, aged between 29 and 74 years) sniffed the samples. Every judge was trained the day before the sample assessment for familiarization with the setup and the odor description procedure. During the GC-O evaluation, judges were voice-recorded and asked to indicate when they perceived an odor (start), its odor quality (descriptors) and when it disappeared (end). The Nasal Impact Frequency (NIF), which is the fraction of judges that detected an odorant at a given time, is plotted as a function of retention time in Fig. 8. A volatile compound was considered odor-active when three or more judges (NIF ≥ 60%) could detect it. Retention Indices (RI) of odor signals were matched with the RIs from the GC-MS data for compound identification.

### Data analysis

To determine the statistical significance between the cell-specific geosmin production in different media, the data was log-transformed and analyzed with one-way ANOVA followed by Tukey’s post hoc test for pairwise comparison. To represent the different volatile profiles of the isolates AT3 and AT4 obtained from the GC-MS analyses, Principal Component Analysis (PCA) on cell-normalized peak areas was performed using MATLAB R2023 (Mathworks, Massachusetts, USA) and the PLS Toolbox (Eigenvector Technologies, Manson, Washington, USA).

## RESULTS

### Identity of two myxobacterial strains isolated from RAS

Two strains that produced cultures with a distinct earthy-muddy scent were successfully isolated. Both strains shared the typical traits of the *Myxococcaceae* family, having slender, rod-shaped vegetative cells with tapered ends, gliding movement in swarms, myxospore and fruiting body formation. Swarms of strain AT3 had a bright green-yellow pigment, as had previously been observed in *Myxococcus virescens* (Lampky & Brockman, 1977; Lang et al., 2008), and fruiting body knob-shaped mounds like that of *Myxococcus* species (Garcia & Müller, 2014). Strain AT4 had hard, densely packed fruiting bodies as previously been observed in *Corallococcus* species (Garcia & Müller, 2014). Indeed, initial 16S rRNA gene sequencing and subsequent phylogenic analysis placed the two isolates AT3 and AT4 within the genera *Myxococcus* and *Corallococcus*, respectively, in the family *Myxococcaceae* (Fig. 1.).

**Fig 1.**
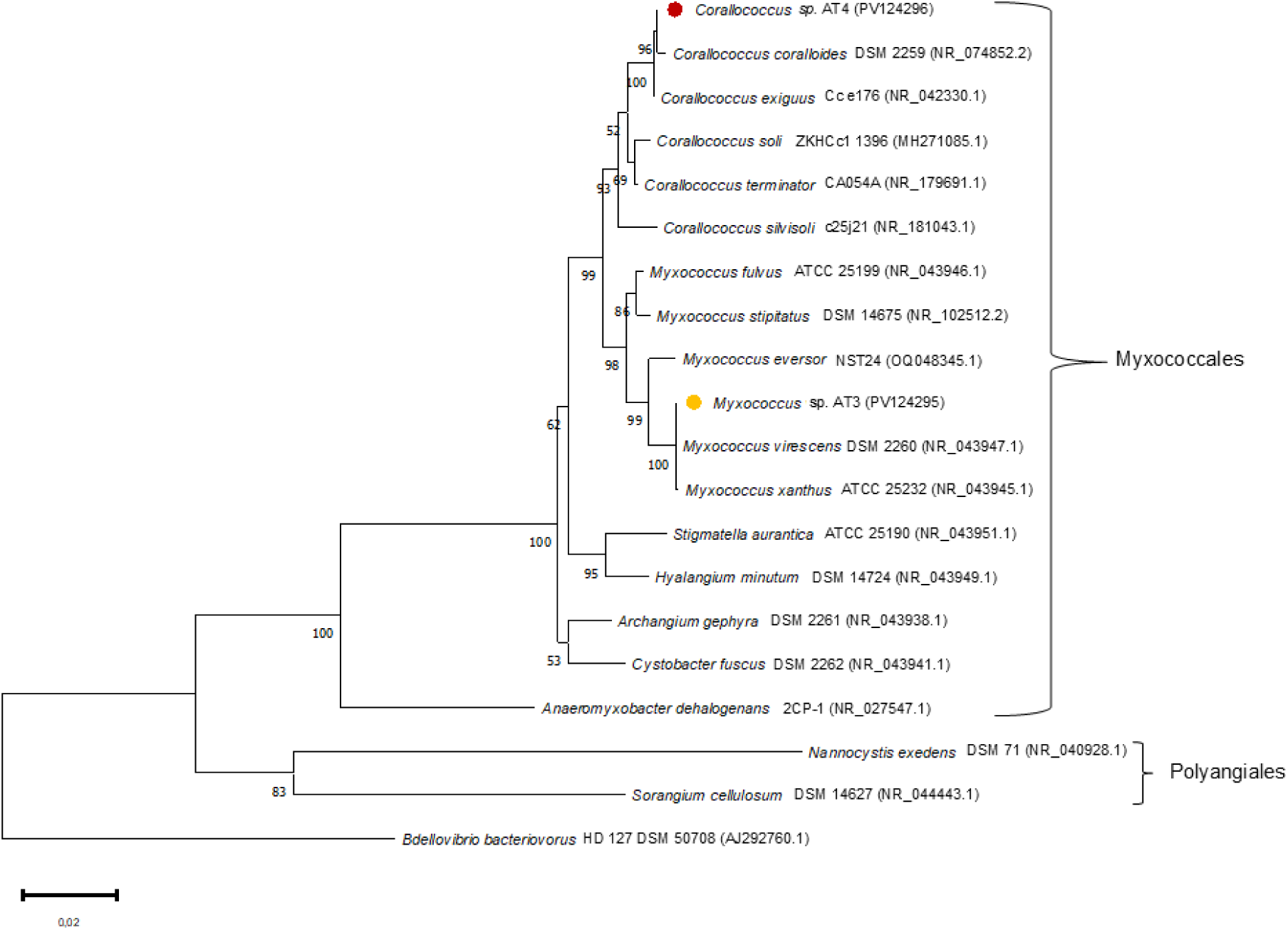
Neighbor-joining tree based on 16S rRNA gene sequences of isolate AT3 (•) and AT4 (•) and related myxobacterial taxa. Accession numbers are shown in parentheses. Bootstrap values (>50%) based on 1000 resamplings are presented above the nodes. The tree is rooted using *Bdellovibrio bacteriovorus* HD127 as an outgroup.

For a deeper taxonomic classification of the isolates, whole genome sequencing was performed. For isolate AT3 the sequencing rendered a 1-contig genome of 9,196,449 bp with 7,467 annotated genes, and a GC-content of 69.1%. Raw sequencing coverage was 94x and assembly coverage 89x. Completeness of the assembly was 99.35%. For isolate AT4 sequencing rendered a 1-contig 10,486,148 bp genome with 8,459 annotated genes and a GC-content of 69.6%. Raw sequencing coverage was 71x and assembly coverage 70x. Completeness of the assembly was 99.35%. Average nucleotide analysis (ANI) revealed that AT3 has genome similarity of 98.85% with *Myxococcus virescens* DSM 2260 and 96.95% of genome similarity with *Myxococcus xanthus* DK 1622 (Fig. 2). Both similarities are above the proposed cutoff value for species delineation with ANI (Goris et al., 2007), however, since AT3 shares phenotypic traits, in particular the green-yellow pigmented swarms, with *M. virescens,* it was determined to belong to this species. AT4 shares 96.40% of its genome with *Corallococcus exiguus* NCCRE002 and is determined to belong to this species. Thus, the isolated strains AT3 and AT4 belong to the species *Myxococcus virescens* and *Corallococcus exiguus*, respectively. Both strains possess the *geoA* gene, as well as a range of secondary metabolite clusters with antibiotic properties, including genes encoding althiomycin, arylomycin, corallopyronin, microsclerodermin, myxalamides and myxochelin, based on antiSMASH analysis.

**Fig. 2.**
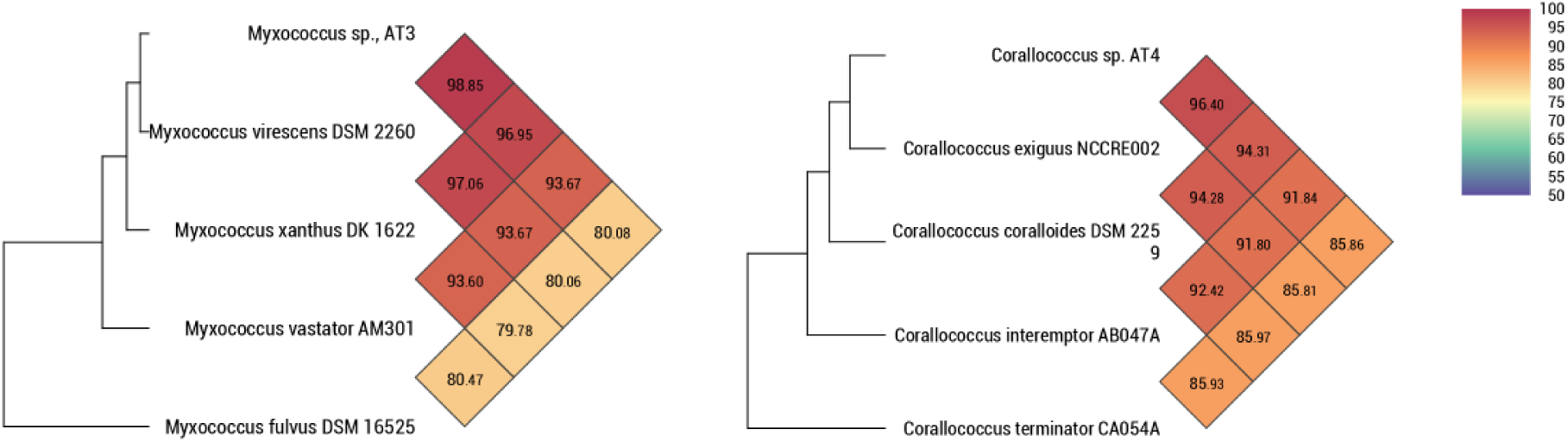
ANI analysis of *Myxococcus* sp. AT3 and *Corallococcus* sp. AT4 with closely related taxa.

### Myxobacteria readily predates on several RAS microbiome members

Members of *Myxococcus* and *Corallococcus* are known to be predatory (Livingstone et al., 2020), and the predatory range of *Myxococcus* on soil bacteria have previously been investigated (Morgan et al., 2010). To determine if strain AT3 and AT4 also have predatory behavior, predation assays using the different bacteria isolated in the RAS biobank were established.

The prey bacteria represented six classes and 11 orders of bacteria. A phylogenetic tree based on the 16S rRNA gene of the prey bacteria, rooted on the two isolated myxobacteria, is shown in Fig. 3. Isolate AT3 and AT4 were capable of preying on 14 of 16 prey bacteria and 15 of 16 of the prey bacteria, respectively. Examples on the predatory behavior is shown for *C. exiguus* AT4 in Fig. 4 Two bacteria, *Pseudomonas* sp. NF1.1 and *Streptomyces* sp. RD2 did initially inhibit the swarming of AT3 and AT4, however, as the incubation progressed, both strains started lysing strain RD2. Only AT4 was capable of lysing strain NF1.1 (Fig. 4C). The excessive swarming of *Flavobacterium onchorynchi* GF28 inhibited the growth of AT4 but not AT3 (Fig. 4B). Neither AT3 nor AT4 lysed *Tahibacter* sp. GF43, nor was their swarming inhibited; instead both myxobacteria just grew closely around the prey (Fig. 4D). Both strains were capable of predation on most bacteria investigated, confirming their broad prey range and potential ecological impact on other RAS bacteria.

**Fig. 3.**
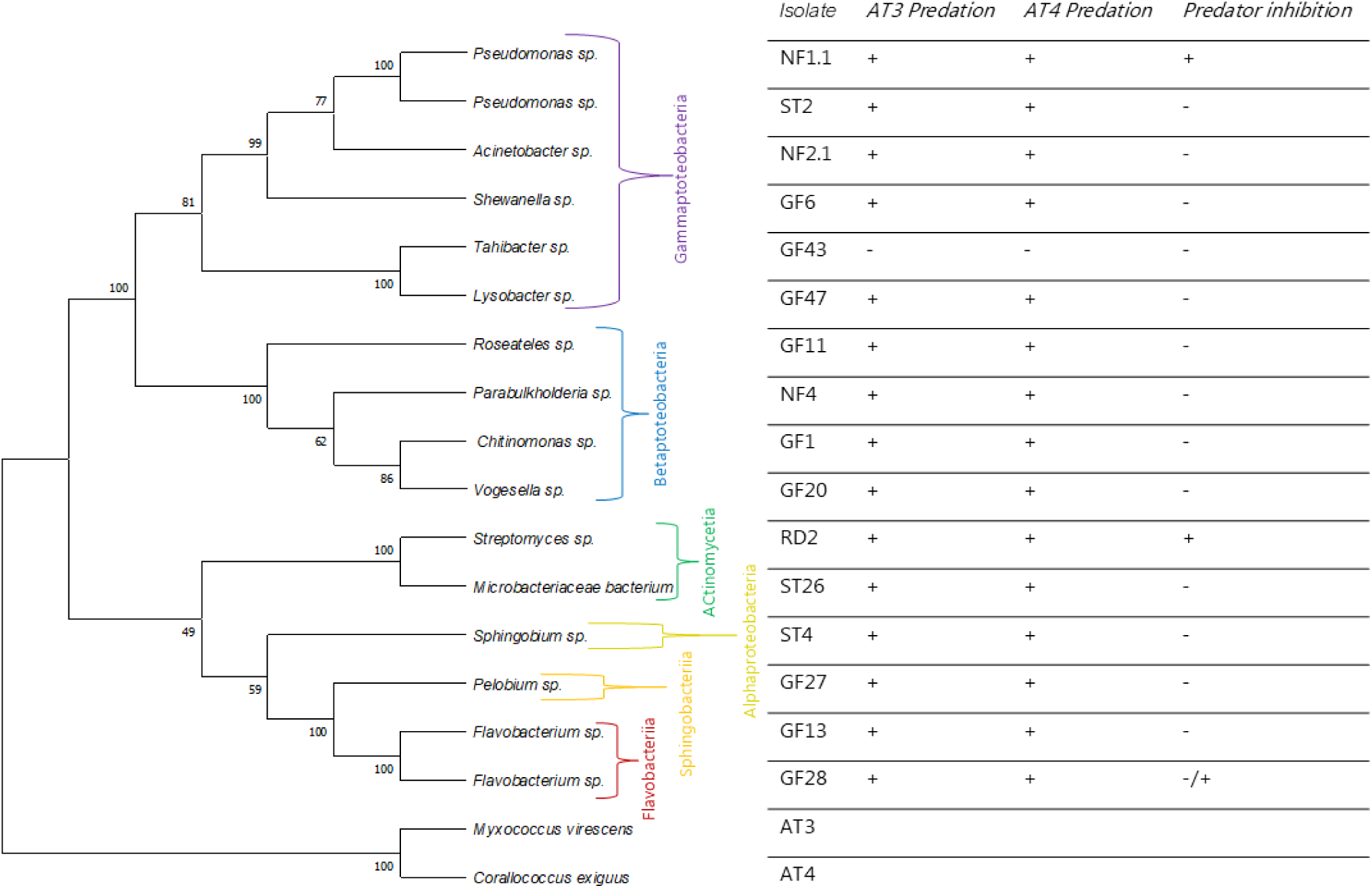
Predation by the myxobacterial isolates AT43 and AT4 on bacteria in the RAS biobank. At left, a neighbor-joining tree based on the 16S rRNA gene sequences of bacteria in the RAS biobank and the two myxobacterial isolates is shown. At right, results from predation by the two myxobacteria on 16 bacterial isolates from the RAS biobank. “+” indicates predation or inhibition of the predator, and “-” indicates the absence of predation or no inhibition of the predator.

**Fig. 4.**
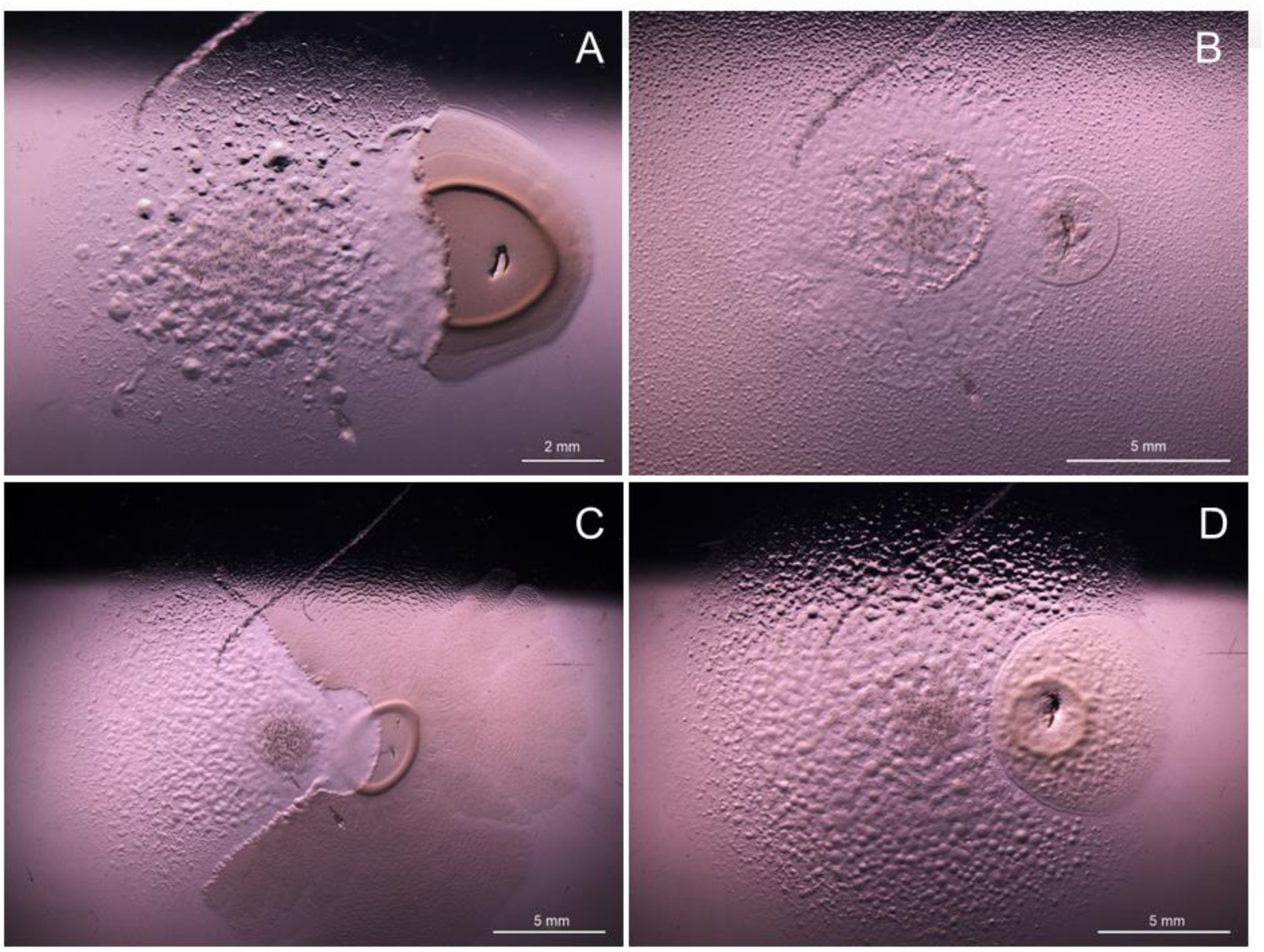
Predator-prey interactions of. (A) *Corallococcus exiguus* AT4 to the left, halfway through the colony of *Acinetobacter* sp. NF2.1 (scale bar is 2 mm); (B) *C. exiguus* AT4 expanding within the swarm of *Flavobacterium* sp. GF28 (scale bar is 5 mm); (C) *C. exiguus* AT4 to the left with swarming *Pseudomonas* sp. NF1.1 (scale bar is 5 mm); (D) *C. exiguus* AT4 growing around a colony of *Tahibacter* sp. GF43 (scale bar is 5 mm).

### Geosmin production depends on nutritional factors

The presence of the *geoA* gene in the two isolates indicated the ability to produce geosmin. To demonstrate geosmin production and possible dependence on the growth conditions of the isolates, AT3 and AT4 were cultivated in different liquid media and geosmin production was quantified. After cultivation, geosmin concentrations were detected and normalized to the number of cells, quantified in each sample by qPCR targeting the 16S rRNA gene and taking into account the different copy numbers of the gene in the respective strains.

The cell-specific geosmin production shows that geosmin was differentially produced by the two strains when cultivated in different media (Fig. 5). The production was significantly higher in the minimal medium and the RAS water than in the other media. A non-significant but inverse trend (AT3 had a high production in RAS water while AT4 had a high production in the minimal medium) was seen between the two strains in these two cultivation settings. Similarly, the cell-specific geosmin production for AT3 was significantly higher in the CTT complex medium and the TPM starvation medium, as compared to the VY/2 complex medium. An inverse relationship between the two strains was found for the VY/2 medium in which AT4 had significantly higher cell-specific geosmin production than in the CTT and TPM media.

**Fig. 5.**
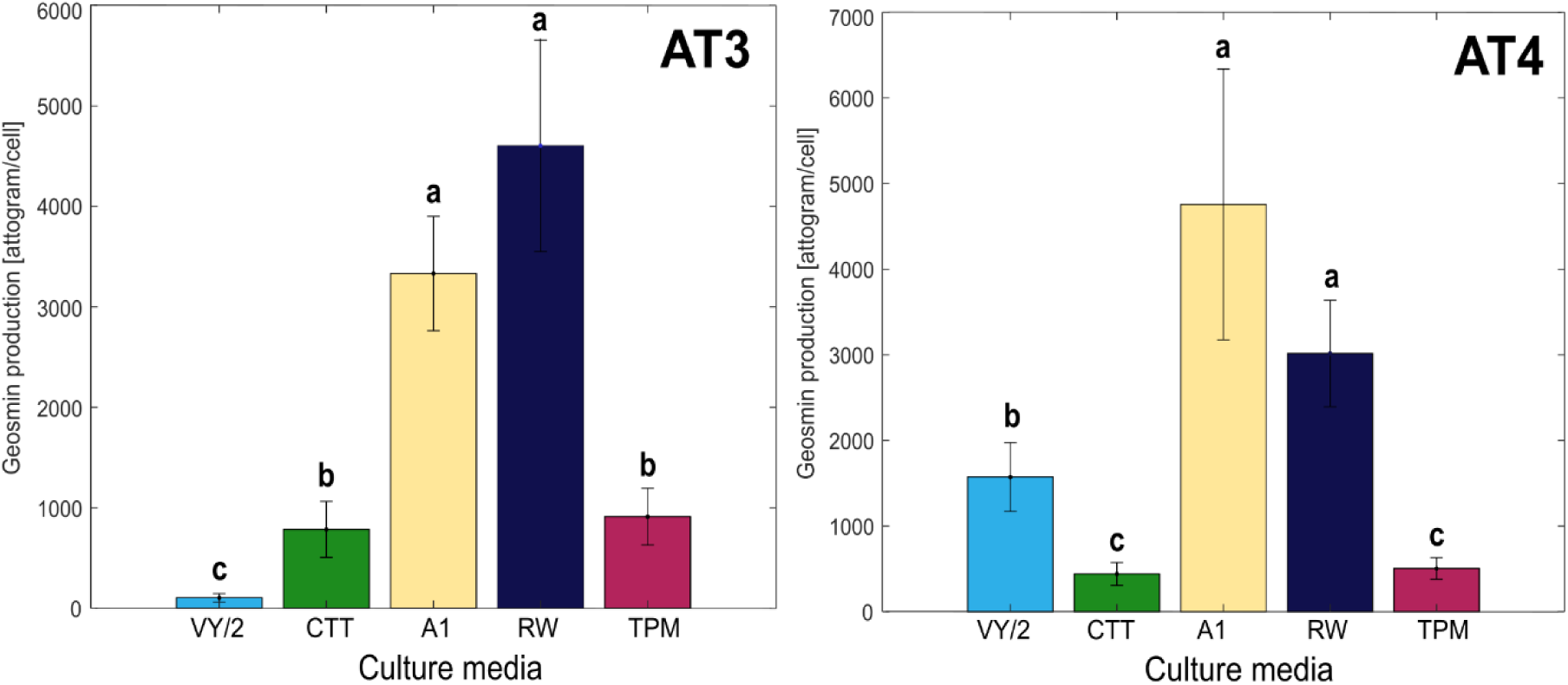
Cell-specific geosmin production of isolate AT3 and AT4 after cultivation in complex media (VY/2 and CTT), minimal medium (A1), RAS rearing water (RW) and starvation medium (TPM). Differences between the samples at P<0.05 level (lowercase letters) and SDs are shown (n = 5).

### Myxobacteria produce other potential off-flavors

Alongside geosmin, strains AT3 and AT4 also produced other VOCs. GC-MS analyses showed that a total of 18 VOCs were produced by the two isolates when cultivated in the different media and when excluding VOCs detected in the control samples (Figs. 6 and 7). Among the 18 VOCs, four were labelled as *unknown* (VOC 3, 4, 11 and 12) because their R.match (spectral similarity value) and Prob (%) (probability of correct identification) values after spectral comparison with the NIST database (2023) were too low (R.match <800 or R match <30%). Two unknown compounds were tentatively identified as mono- and sesquiterpenes (VOC 17 and 18), given the spectral similarity with compounds from these terpene classes (Appendix A1). PCA biplots of the volatile profiles (normalized to the number of cells) from the cultivation of AT3 and AT4 in the different media are shown in Figs. 6 and 7.

**Fig. 6.**
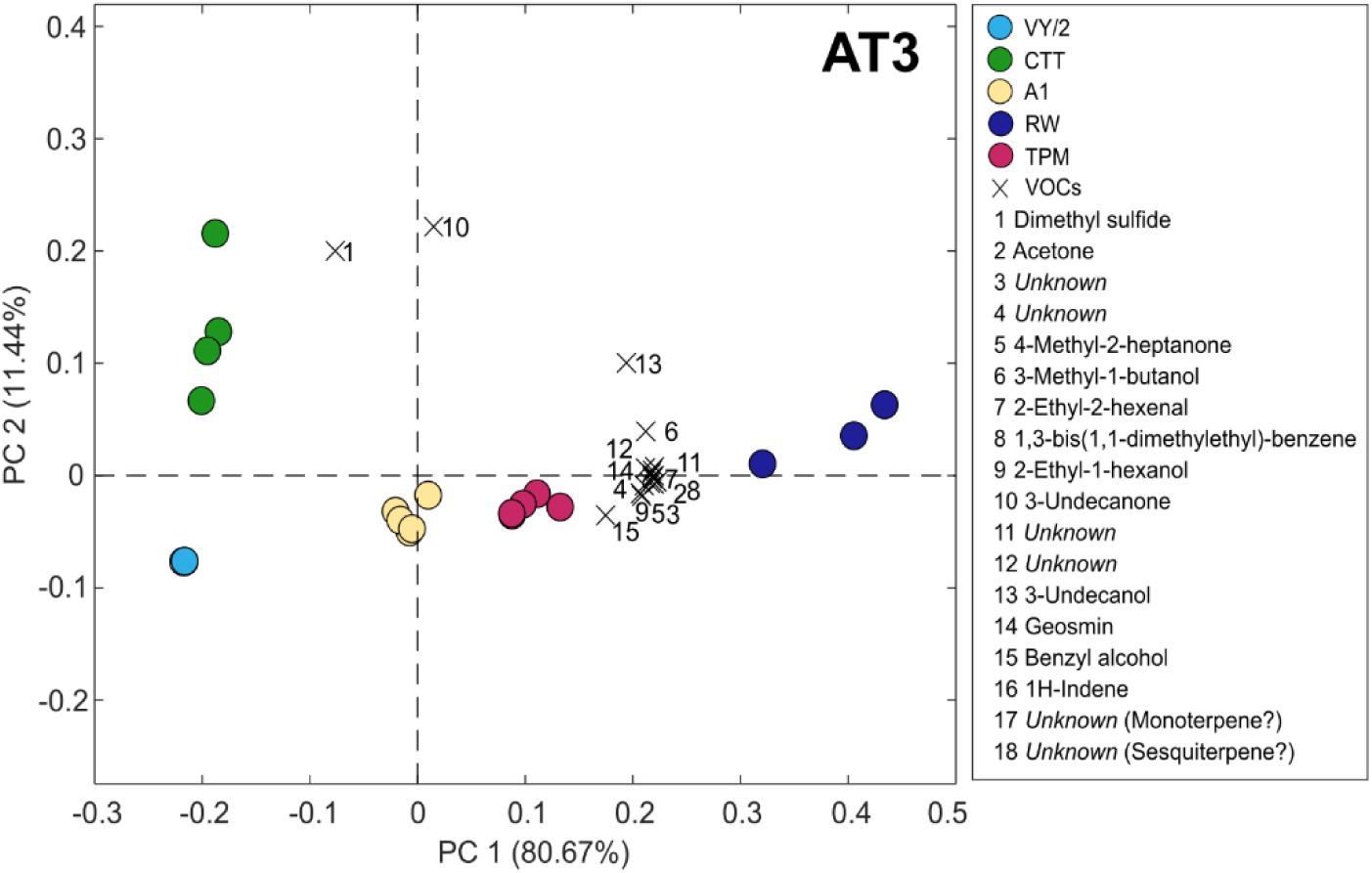
PCA biplot of VOCs produced by AT3. Identified compounds are designated number 1 to 18. Colored circles represent the number of samples cultivated in different media.

**Fig. 7.**
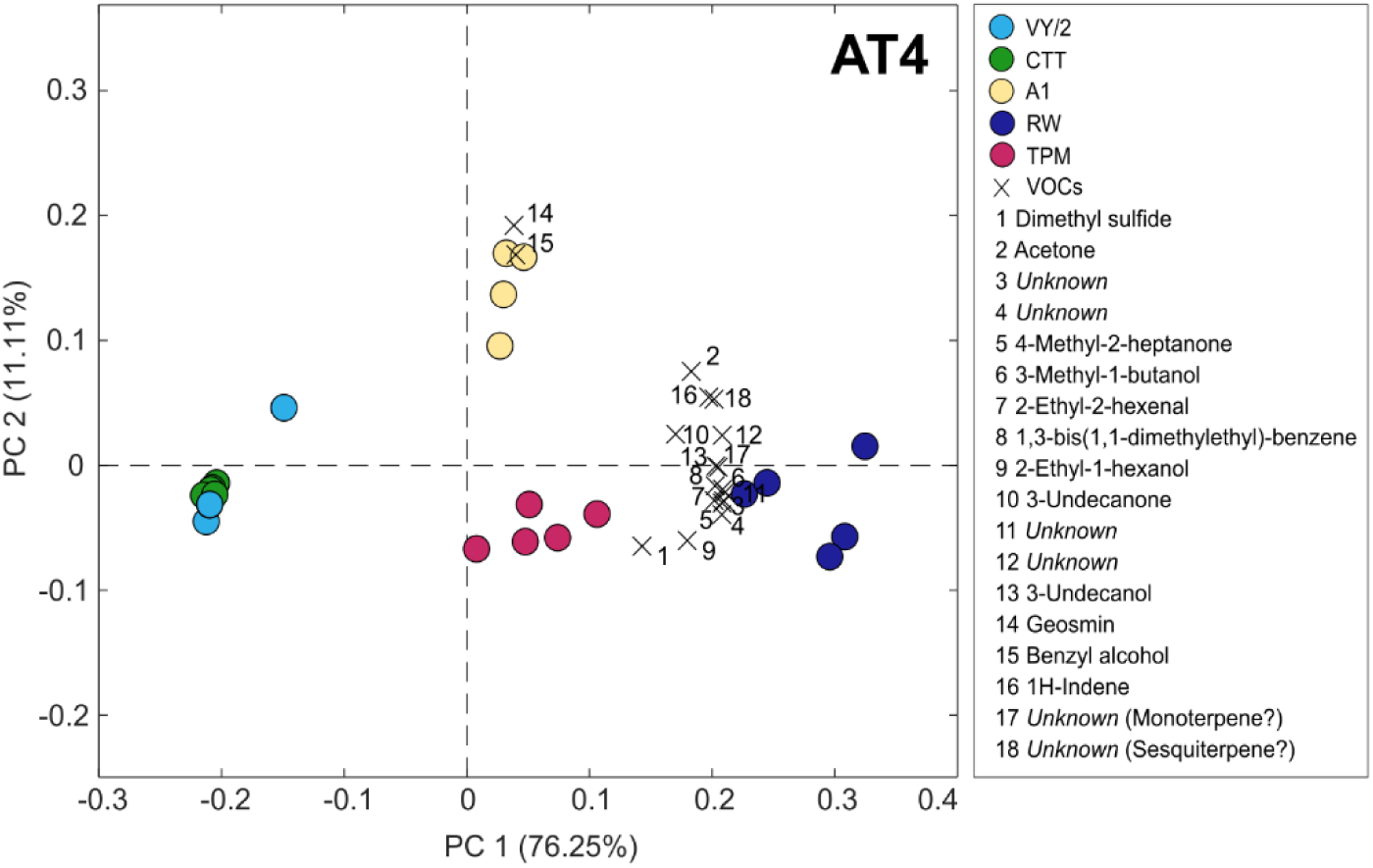
PCA biplot of VOCs produced by AT4. Identified compounds are designated number 1 to 18. Colored circles represent the number of samples cultivated in different media.

For AT3, PC1 in the biplots explains 80.67% of the variance and clusters the samples along PC1, while PC2 explains 11.44% of the variance (Fig. 6). The most separated samples with respect to PC2 occurred from cultivation in CTT and VY/2 media, as shown in quadrants III and IV. Especially cultivation in RW but also in TPM was associated with a higher production of most of the detected VOCs. The exceptions were dimethyl sulfide (1) and 3-undecanone (10), which showed no strong associations with cultivation in any of the different media except for CTT.

For AT4, PC1 also captures most of the systematic variance in the data (76.25%) and distinctively clusters the samples along PC1. PC2 explains 11.11% of the variance (Fig. 7). In this case, PC2 does not separate the cultivation on CTT and VY/2 but explains the higher production of geosmin (14) and benzyl alcohol (15) associated with A1. In the RW medium, the samples were associated with a higher production of most VOCs (except for 14 and 15).

In addition to identifying the VOCs produced by the two strains, the sensory relevance of these compounds in terms of odor attributes and intensity is necessary information to determine their potential influence in off-flavor generation. This is especially relevant for the two unknown presumptive terpenoids (VOC 17 and 18), since many terpenoids have characteristic aroma profiles. For this purpose, gas chromatography-olfactometry (GC-O) was employed as a screening tool to discriminate the odor-active compounds produced. Strain AT3 cultivated in CTT medium was selected for this analysis, as the absolute production of volatiles was highest on this medium, maximizing the extraction and detection of VOCs with potential odor activity. After the evaluation sessions, an aromagram was constructed with 12 identified odors, based on the established critera (NIF ≥ 60%) (Fig. 8A). The matching of these odorants by odor description and retention index (RI) and identification by GC-MS is shown in Table 2. The analysis helped us distinguish between the smelled compounds belonging to the CTT medium and the volatiles that exclusively were produced by AT3. Out of the 12 odors registered in the GC-O analysis, only five could be identified in the GC-MS analysis of VOCs produced by AT3 (VOCs produced by AT3 are indicated in green color in Fig. 8B). These five odors were dimethyl sulfide (1), 4-methyl-2-heptanone (5), 3-methyl-1-butanol (6), geosmin (14) and an unknown compound (18), which has the electron ionization mass spectrum (EI-MS) of a sesquiterpenoid. The remaining VOCs were assumed originating from the medium (red color in Fig. 8B) or were unknowns (black color in Fig. 8B).

**Table 2.** Odorants detected by GC-O of AT3 cultivated in CTT medium. Odor descriptions, associated chemical compound, RI and source of origin are specified.

| Odor # | Odor descriptions | Compound | RI<sub>GC-O</sub> | RI<sub>GC-MS</sub> | RI<sub>standard</sub> | Source |
| --- | --- | --- | --- | --- | --- | --- |
| Odor 1 | Sulfury, green | Dimethyl sulfide | 735 | 745 | - | AT3 |
| Odor 2 | Earthy, nutty | 1-Butanol | 1133 | 1165 | 1165 | CTT |
| Odor 3 | Forest, organic acid | 4-Methyl-2-heptanone | 1211 | 1212 | - | AT3 |
| Odor 4 | Medicinal, chemical | 3-Methyl-1-butanol | 1224 | 1224 | 1224 | AT3 |
| Odor 5 | Floral, metallic, hops, flower | 2,5-Dimethylpyrazine | 1337 | 1338 | 1335 | CTT |
| Odor 6 | Sulfury | Nonanal | 1407 | 1401 | 1404 | CTT |
| Odor 7 | Musty, green bell pepper, nutty | Pyrazine? | 1419 | 1417 | - | CTT |
| Odor 8 | Green bell pepper | 3-Ethyl-2,5-dimethyl-pyrazine | 1456 | 1459 | - | CTT |
| Odor 9 | Green bell pepper, earthy, fruit | Unknown* | 1560 | - | 1542 | Unknown |
| Odor 10 | Musty, earthy, leather | Unknown** | 1573 | - | 1551 | Unknown |
| Odor 11 | Geosmin | Geosmin | 1900 | 1856 | 1854 | AT3 |
| Odor 12 | Musty, earthy, flowery | Sesquiterpene? | 2232 | 2176 | - | AT3 |

\*Tentative identification of 3-isobutyl-2-methoxypyrazine/2-sec-butyl-3-methoxypyrazine by odor description and RI of an authentic standard. No MS from the sample is available.
\*\*Tentative identification of (E)-2-nonenal by odor description and RI of an authentic standard. No MS from the sample is available.

**Fig. 8.**
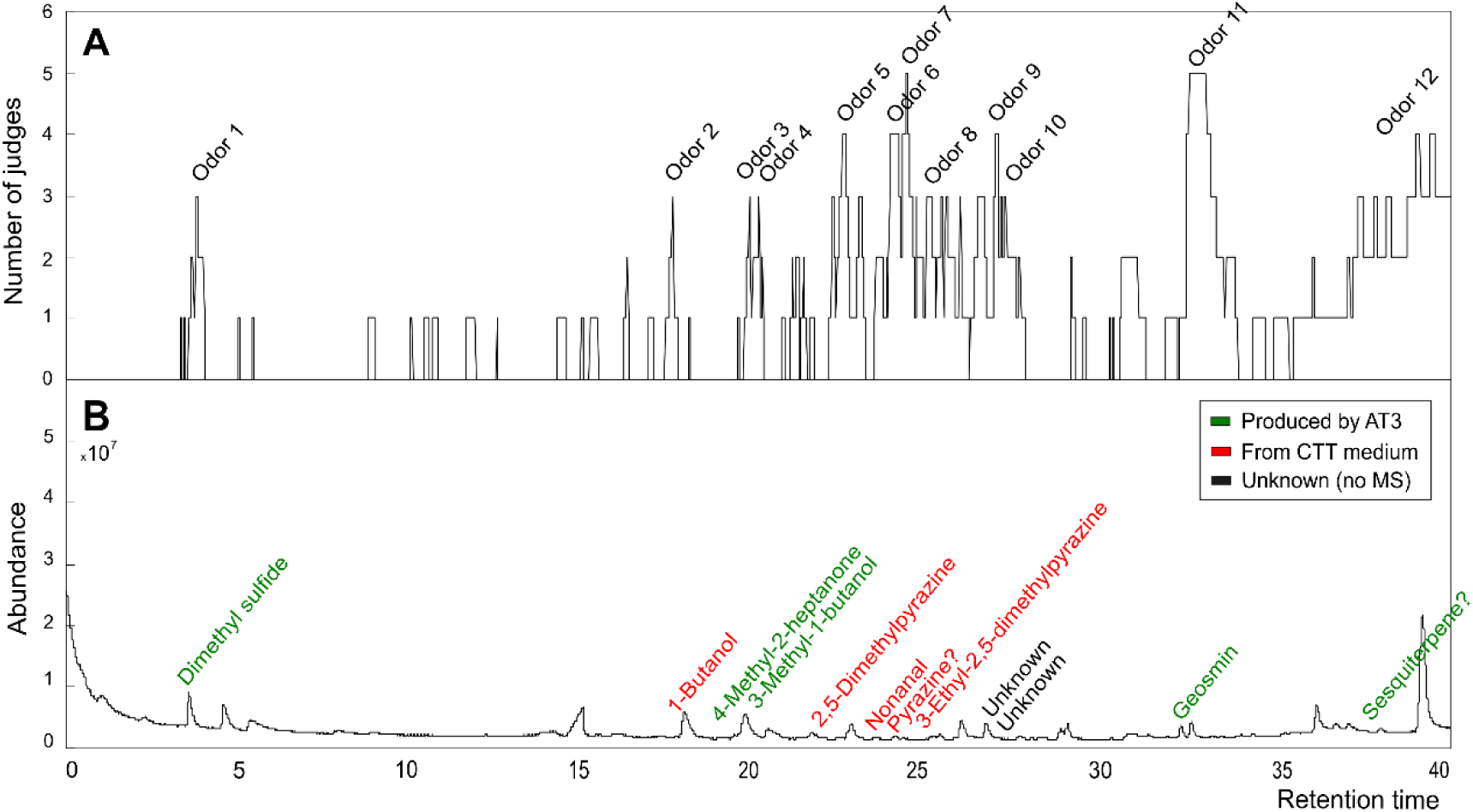
(A) Aromagram of AT3 cultivated in CTT assessed by five judges. Twelve odors had nasal impact frequency (NIF) equal to or above 60%. (B) GC-MS chromatogram of AT3 cultivated in CTT displaying the compounds that were found and that matched the odors perceived. Green color shows the compounds that were produced by AT3, while red color shows compounds associated with the culture media, and black color shows the unknown compounds linked to two specific odor regions.

## DISCUSSION

The presence of potentially geosmin-producing myxobacteria in RAS has previously been documented through molecular methods (Auffret et al., 2013; Lukassen et al., 2017, 2022) but none of the geosmin-producing strains were isolated. In our study, we successfully isolated two *geoA*-harboring myxobacteria from RAS, identified as *Myxococcus virescens* AT3 and *Corallococcus exiguus* AT4. Their geosmin production has been confirmed and quantified via GC-MS analysis, and their production of other potential off-flavor compounds was investigated via GC-O.

### Geosmin production vs. nutrient availability

Strains of *Myxococcus* and *Corallococcus* are among the most frequently isolated myxobacteria (Garcia & Müller, 2014). The two isolates in our study were identified to belong to the species *Myxococcus virescens* and *Corallococcus exiguus*. These two species have primarily been found in soil (Microbe Atlas, 2025). In RAS environments, previous studies (qPCR of the *geoA* gene) indicated that members of the genus *Sorangium* were the most abundant geosmin-producing myxobacteria (Lukassen et al., 2017), but no information on geosmin production was provided. In the present study, the geosmin production of the two myxobacterial strains reached average levels of 100-4800 ag (ag = 10^-18^ gram) geosmin per cell after two days of cultivation. To our knowledge, there is only one other published study in which geosmin production by myxobacteria was quantified. Yamamoto and coworkers determined the geosmin production in several *Myxococcus* isolates from water and lake sediment, cultivated in CY medium, and measured rates of 8-270 µg/L after a three-day incubation period, but the cell-specific rates were not presented (Yamamoto et al., 1994). If comparing the cell-specific geosmin production in our study to that of *Streptomyces* isolated from Danish fish ponds, determined to 0.1 to 35 ag geosmin bacterium^−1^ h^−1^ across the six *Streptomyces* isolates, the production in the present study ranges from 2 to 99 ag geosmin bacterium^−1^ h^−1^ when calculated as in Klausen et al. (2005). However, it should be noted that Klausen et al. only carried out cultivation in a nutrient-rich medium. In our study, the nutrient-rich media resulted in the lowest cell-specific production rates, although an inverse relationship for the two strains was found for cultivation in the VY/2 and CTT media, and the A1 minimal medium and the RAS water medium led to the highest rates. The different rates indicate that geosmin production in myxobacteria is not constitutive. From a RAS point of view, this could be considered a positive finding as it indicates that, at least in theory, the geosmin production by these bacteria can be manipulated.

The applied A1 minimal medium was developed for the model myxobacterium *M. xanthus* and is supposed to provide the lowest amounts of nutrients required to support growth in this bacterium (Bretscher & Kaiser, 1978). In the A1 medium, the generation time was determined to range from 22 to 30 h under optimal temperature, while the CTT medium, which has a considerably higher nutrient content, reduces the generation time of *M. xanthus* to 4-6 h (Schumacher & Søgaard-Andersen, 2018)Klik eller tryk her for at skrive tekst.. The exact composition of the presently applied RAS water (RW) medium at the time of sampling is unknown, but nutrient levels in rearing water from this farm have previously been found to 9-15 mg mg/L nitrate, 0.50-0.85 mg mg/L ammonium, and 1.8-321 µg/L phosphate during a three-month period (Podduturi et al., 2025).

A relationship between geosmin production and nutrient availability seems evident by the significantly higher production in the low-nutrient medium A1 and the RAS rearing water (RW), as compared to the complex media that better supports the growth of both strains. A similar pattern of higher geosmin production during suboptimal growth conditions has been observed in Cyanobacteria. High nitrate concentrations stimulated the growth but decreased the geosmin production in *Dolichospermum smithii* (Shen et al., 2021a). A related trend was observed for *Anabaena viguieri* under high ammonia concentrations, which increased growth but lowered the geosmin production (Wu et al., 1991). While cyanobacterial growth is stimulated by light and optimum temperature, the highest geosmin production has been shown to occur at suboptimal light and temperature (Shen et al., 2021b; Wang & Li, 2015; Zhang et al., 2009, 2017). It has been theorized that since chlorophyll and geosmin production both depend on the terpenoid biosynthetic pathways, geosmin production is down-prioritized for chlorophyll production, when conditions are optimal for growth (Wang & Li, 2015).

Inorganic nutrients have also been found to affect the geosmin production in *Streptomyces*, although the results were variable. When *Streptomyces* spp. strain S10 was cultivated in basal salt medium with different concentrations of phosphate and nitrate, it had the highest geosmin concentration during low phosphate concentrations (0.05 mg/L) and high nitrate concentrations (100 mg/L) (Shudirman et al., 2021). These concentrations are considerably higher than those expected in a RAS setting (Davidson et al., 2017) and may not relevant for RAS farming. Interestingly, in contrast to geosmin, 2-MIB was only produced by this *Streptomyces* under low nitrate conditions (Shudirman et al., 2021). In another study of *Streptomyces*, *S. halstedii,* the highest geosmin production was determined under low ammonia and nitrate (Blevins et al., 1995). Low levels of the micronutrients iron and copper have also been found to stimulate geosmin production in *S. halstedii* (Schrader & Blevins, 2001).

A common characteristic of the A1 and RW media is the low nutrient content compared to the two complex media VY/2 and CTT. While levels of available carbon might influence geosmin production, no such correlation was found in a recent study comparing water quality parameters and geosmin in the water in a commercial RAS (Podduturi et al., 2025). It is uncertain whether the general observation of increased geosmin production under low nutrient availability is caused by similar biochemical or biological mechanisms between groups that are phylogenetically and biologically very distant. A speculation might be that geosmin acts as an intercellular signal to initiate movements involved in myxobacterial swarming and in the movement of cyanobacterial filaments and streptomycete hyphae, and possibly also in cellular differentiation (Bush et al., 2022; Muñoz-Dorado et al., 2016; Schuergers et al., 2017). In our study, cultivation of the two isolated myxobacteria in TPM (starvation medium known to induce sporulation (Caberoy et al., 2003)), did not generate a higher geosmin production than the two low-nutrient media. This suggests that geosmin production is not substantially upregulated during sporulation as found in *Streptomyces* (Becher et al., 2020).

### The contribution of other VOCs to off-flavor

VOC production in microorganisms is influenced by the nutritional conditions of their environment (Weisskopf et al., 2021), as well as by biosynthetic pathways (primary metabolism, fermentation, terpene pathway, etc.), resulting in many different chemical structures (Schulz & Dickschat, 2007; Weisskopf et al., 2021). In the present study, the GC-MS analysis indicated that a total of 18 volatile compounds were produced by *M. virescens* AT3 and *C. exiguus* AT4. The GC-O analysis indicated that additional VOCs occurred in the cultures, possibly originating from the culture medium (Fig. 8B). Speculatively, these compounds could also be produced by the bacterial strains. For example, 2,5-dimethylpyrazine is a rather widespread VOC across bacterial groups (Schulz & Dickschat, 2007) and 3-ethyl-2,5-dimethylpyrazine was identified as a VOC from North Sea bacteria (Dickschat et al., 2005B).

The cultivation of AT3 and AT4 in RAS water indicated a stimulated production of several volatiles and not exclusively geosmin (Figs. 6 & 7). Yet, the GC-O analysis showed that geosmin was undoubtedly one of the major odor-active compounds (NIF = 100%, detected by all panellists). This reaffirms the potency of geosmin and its relevance in contributing earthy-musty flavor notes in RAS-reared fish. Other compounds of particular interest were two unknown compounds (VOC 17 and 18) with mass spectra closely resembling those of certain monoterpenes and sesquiterpenes (Appendix A1). In the GC-O analysis of *M. virescens* AT3 cultivated in the CTT medium, the likely monoterpene (17) could not be detected, but the presumptive sesquiterpene (18) was described by the assessors as having “musty”, “earthy” and “flowery” attributes, hence being another potential compound contributing earthy-musty taste in RAS-farmed fish.

The volatile compounds benzyl alcohol, 3-undecanone and 3-undecanol produced by AT3 and AT4 (Figs. 6 & 7) have previously been reported to be produced by other myxobacterial strains (Dickschat et al., 2005; Schulz et al., 2004; Xu et al., 2011) but they were not odor-active according to the GC-O results. Acetone, 2-ethyl-2-hexenal, 1,3-*bis*-(1,1-dimethylethyl)-benzene and 1H-indene are here reported for the first time as VOC produced by myxobacteria, but none of them was perceived as odor-active by the panelists. The compound 2-ethyl-1-hexanol was also identified in this study and has been reported as a microbial degradation product of plasticizers. This means that it can be deemed as an artefact VOC that does not originate from the strain nor the media, but most likely came from the cultivation tubes (Nalli et al., 2006; Schulz & Dickschat, 2007). The compounds detected by the GC-O analysis and originating from strain AT3 (green color in Fig. 8) were dimethyl sulfide (described as “sulfury” and “green”), 4-methyl-2-heptanone (described as “forest” and “organic acid”); and 3-methyl-1-butanol (described as “medicinal” and “chemical”). Dimethyl sulfide is produced by a range of microorganisms (Schulz & Dickschat, 2007), perhaps the most well-known source is marine phytoplankton (Cuong Nguyen et al., 1988), and it has been reported as an off-flavor-causing compound in fish feed and rainbow trout flesh (Noguera et al., 2024). The compounds 4-methyl-2-heptanone and 3-methyl-1-butanol have not previously been reported as off-flavors in RAS facilities, nonetheless they were both described with disagreeable odor qualities. The production of 3-methyl-1-butanol has been previously reported in the myxobacterium *S. cellulosum* (Xu et al., 2011). Finally, odor 9 (described as “green bell pepper”, “earthy”, “fruit”) and odor 10 (described as “musty”, “earthy”, “leather”) are tentatively identified as a 3-isobutyl-2-methoxypyrazine/3-sec-butyl-3-methoxypyrazine and (*E*)-2-nonenal. The odor descriptions provided by the panellists match their reported odor qualities (Kreissl et al., 2022), as well as their linear retention indices, which also matched the retention indices of the authentic standards. However, no mass spectra of these compounds could be determined in the myxobacteria samples. Both compounds have extremely low odor thresholds (in the parts-per-trillion range) (Sidhu et al., 2015), which is likely the reason why they were not detected by GC-MS analysis but still be perceived by the panellists. Further analyses are needed to confirm the production of these relevant odor-active compounds by AT3 and AT4.

### Predator-prey interactions between myxobacteria and other RAS bacteria

Myxobacteria require surfaces to maintain their multicellular lifestyle (Pathak et al., 2012; Wrótniak-Drzewiecka et al., 2016). In a RAS environment, they presumably inhabit biofilter media, compartment walls, and solid particles. As they are both saprophytic and predatory, their diet in RAS probably consists of organic matter generated in the system, as well as preying on other members of the RAS microbiome. In the predation assay on 17 bacteria isolated from four different RAS, both *M. virescens* AT3 and *C. exiguus* AT4 could readily predate on most of the bacteria tested, demonstrating a large predatory potential. This agrees with previous reports on these two genera (Inoue et al., 2022; Livingstone et al., 2020). Surprisingly, only *C. exiguus* had a successful predation on *Pseudomonas* sp. NF1.1. This may show the ability of *Pseudomonas* to defend itself from predation, as has been observed previously, probably reflecting defense mechanisms involving antibiotic resistance through efflux pumps, mucoid conversion, and formaldehyde excretion (Akbar & Stevens, 2021; Sutton et al., 2019). The only bacterium that completely avoided predation by both myxobacterial strains was *Tahibacter* sp. GF43. In the predation assay, it became closely surrounded by both AT3 and AT4 but without being lysed. It is unknown which mechanisms caused this predation avoidance, as there are few studies on this bacterium. However, it might be speculated that excretion of extracellular polymeric substances protects *Tahibacter* from predation in a similar way as has been observed with avoidance of predation of *Sinorhizobium meliloti* from *M. xanthus* (Pérez et al., 2014).

## Conclusion

Myxobacteria have generally been considered as mainly soil bacteria (Dawid, 2000). Although their proportion of prokaryotic populations in soils probably is less than 4.5%, predatory myxobacteria have been proposed to be keystone taxa in soil microbial food webs, assumed to significantly influence the structure of soil microbial communities due to their predation (Li et al., 2024; Petters et al., 2021). The present identification and isolation of myxobacteria in fish farm water shows that these bacteria also occur in aquatic environments as previously suggested in molecular analysis (Lukassen et al., 2022).

In this study, myxobacteria have, for the first time, been isolated from a RAS environment, enabling in-depth studies of their production of geosmin and other potential compounds contributing to earthy-muddy tasting fish. Quantification of their geosmin production shows that they are prolific geosmin producers, comparable to previously quantified levels of *Streptomyces* cultures. The geosmin production seems to be stimulated by cultivation in low-nutrient media, including the cultivation in rearing water from RAS. This might indicate that the comparatively low-nutrient environment in RAS upregulates the geosmin production in these bacteria. Apart from geosmin, the strains produced other previously known and novel, unknown off-flavor compounds, underlining the importance of further investigation of the influence of these bacteria and their metabolites on off-flavors in RAS production. The assays on the predatory potential of the two myxobacterial strains, targeting also several RAS-native taxa, demonstrate their presumable ecological impact on the RAS microbiome. This study emphasizes that myxobacteria due to their predatory and metabolic potential are important components of the microbial ecology in RAS facilities.

## ACKNOWLEDGEMENTS

The present study was funded by the European Union’s Horizon and innovation programme under the Marie Skłodowska-Curie grant agreement No. 956481 (RASOPTA).

## AUTHOR CONTRIBUTION

**Julia Södergren,** Writing – original draft, Conceptualization, Methodology, Investigation, Formal analysis, Visualization I **Pedro Martínez Noguera,** Writing – original draft, Methodology, Investigation, Formal analysis, Visualization I **Mikael Agerlin Petersen,** Writing – review and editing, Methodology, Investigation, Formal analysis I **Niels O. G. Jørgensen,** Writing – review and editing, Investigation, Conceptualization, Supervision, Funding acquisition I **Raju Podduturi,** Writing – review and editing, Conceptualization, Supervision I **Mette H. Nicolaisen,** Writing – review and editing, Methodology, Investigation, Conceptualization, Supervision.

## DATA AVAILABILITY

Data will be made available on request.

## Appendix A1

### EI-MS Fragmentation pattern of VOC 17

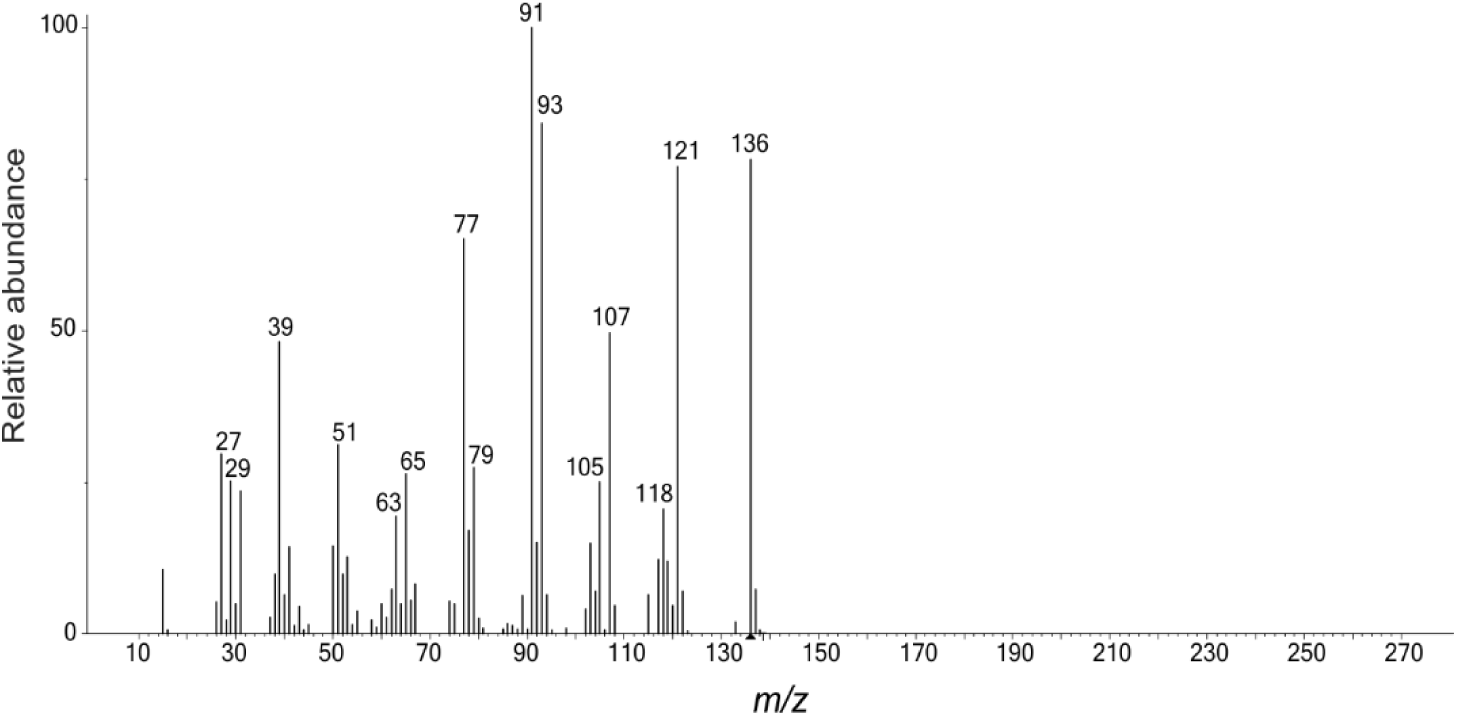

Monoterpene hydrocarbons, regardless of whether they are acyclic, monocyclic or bicyclic, typically yield fragmentation patterns under EI-MS conditions like the MS acquired from VOC 17. *m/z* 136 (C_10_H_16_) corresponds to the molecular ion (M), while *m/z* 121 (C_9_H_13_) and *m/z* 93 (C_7_H_9_) correspond to M-15 and M-43 fragments. However, the relative signal intensities of the different ions vary across compounds and would therefore be difficult to pinpoint which monoterpene class VOC 17 belongs to. Among monoterpene alcohols produced by another myxobacterium, *Stigmatella aurantiaca*, were menthol, p-menth-1-en-4-ol and α-terpineol, but no monoterpene hydrocarbons have been reported so far in myxobacteria (Schulz & Dickschat, 2007).

### EI-MS Fragmentation pattern of VOC 18

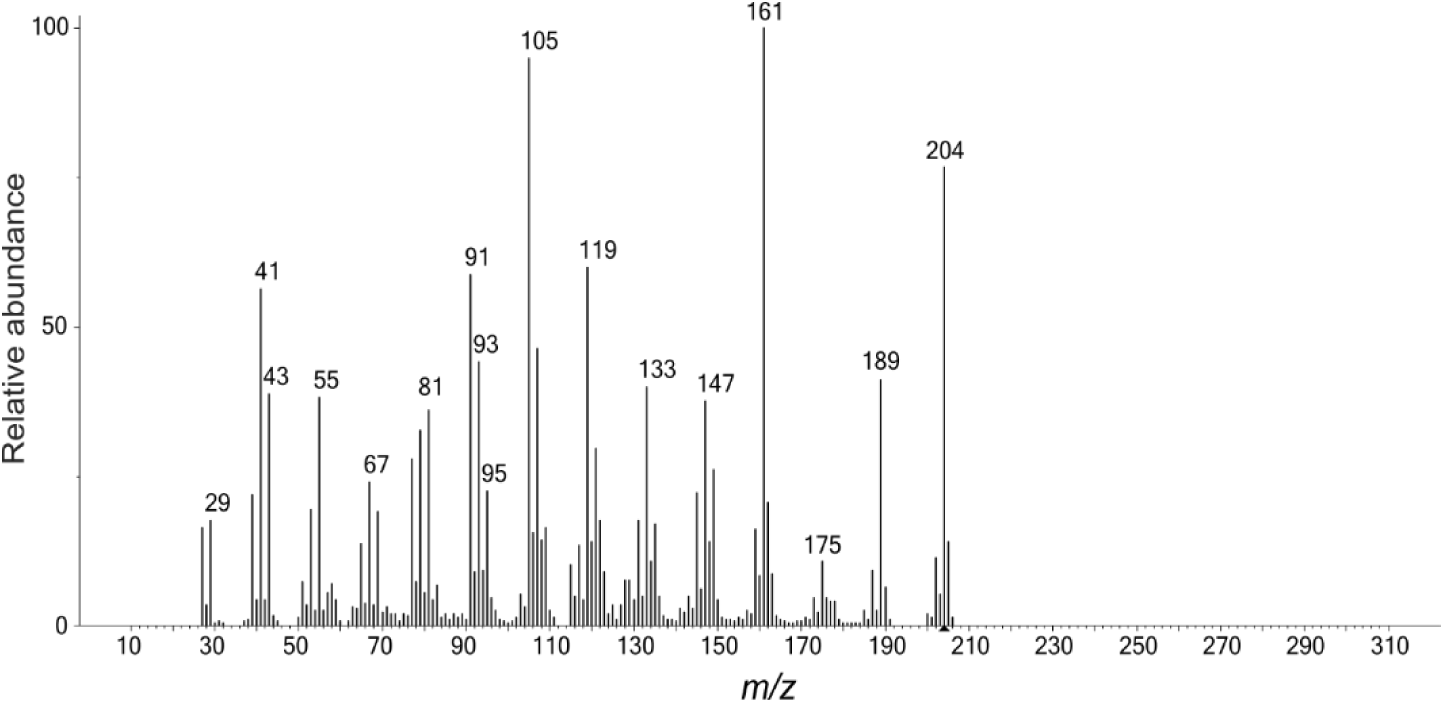

Sesquiterpenes, such as longifolene, cadinene or aromadendrene, are reported to have an EI-MS fragmentation pattern comparable to the one found for VOC 18. *m/z* 204 (C_15_H_24_) corresponds to the molecular ion (M) and *m/z* 189 (C_14_H_21_) and *m/z* 161 (C_12_H_17_) to typical M-15 and M-43 fragments. As for monoterpenes, the abundances of these ions vary across sesquiterpenes. An array of sesquiterpenes have been reported as produced by different myxobacterial strains. For instance, β-ylangene, β-copaene, *iso*-germacrene-D, (-)-germacrene D (also found in *M. xanthus* (Dickschat et al., 2004)), eremophilene, zonarene were identified in the headspace of agar plate cultures of *C. crocatus* and *S. aurantiaca* (Dickschat et al., 2005A; Schulz et al., 2004).

